# Comparative transcriptomics of THP-1 monocytes in response to different pathogens

**DOI:** 10.1101/155853

**Authors:** Ernst Thuer, Toni Gabaldón

**Affiliations:** Centre for Genomic Regulation (CRG), The Barcelona Institute of Science and Technology, Dr. Aiguader 88, Barcelona 08003, Spain; Universitat Pompeu Fabra (UPF). 08003 Barcelona, Spain.; ICREA, Pg. Lluís Companys 23, 08010 Barcelona, Spain.

## Abstract

Undifferentiated human monocytes encounter various pathogens while present in the bloodstream. They are considered a primary responder and regulator for human immune reactions. As such, experiments investigating responses to pathogens are often reliant on monocyte cell cultures. For reproducibility reasons, immortalized cell lines are used. One of the most important cell lines used to model pathogen interactions is THP-1, which has been used in a variety of high throughput transcriptomics experiments. Yet as a cancer derived cell line it may no longer maintain its orignial functionality in detecting and responding to pathogens. Using available large scale transcriptomics datasets, we compare the response of THP-1 to a variety of human pathogens; viruses, bacteria, protozoa and fungi. Our approach focuses on the behavior of THP1 in its response to the different pathogens. Our aim is to provide comparative insights into the cell lines, which may serve to potentially improve future experimental design.

## 1 Introduction

Undifferentiated human monocytes reside in the bloodstream for up to three days before differentiating and moving into tissue. During this time, they are a primary responder to any invasive pathogen entering the bloodstream. Due to their primary role in coordinating host responses, they have been suggested as targets for immune augmentations strategies e.g during fungal infections (Segal, 2007). Most experiments investigating bloodstream infections to pathogens rely on the use of immortalized cell lines to reliably and reproducibly model potential interactions between humans and pathogens e.g (Leland and Ginocchio, 2007). A potential downside of such cell lines, usually derived from cancer lines, is that they are established after human cells have already mutated to exhibit unnatural behaviour (Kaur and Dufour, 2012).

A common cell line used to study the behavior of monocytes is the leukemia derived cell line THP-1 (Tsuchiya et al., 1980). This cell line has been used to study a variety of pathogens using RNA sequencing based transcriptomics. Experiments using THP-1 interaction models involve interactions with viruses, Ebola and Marburgvirus (Martinez et al., 2013), Zika (Hanners et al., 2016) and bacteria such as *Coxiella burnetii* (Millar et al., 2015), *Mycobacteria spp*. (Reyes et al., 1999; Zakharova et al., 2010) as well as the protist *Leishmania mexicana* (Millar et al., 2015). In a recent study, Toth et al. (Tóth et al., 2017) Investigated the response of the pathogenic yeast *Candida parapsilosis* to THP-1. Additionally, available data includes exposure to ethanol, calcitriol and Tissue-type Plasminogen Activator (TPA) (Barendsen, Mueller, and Chen, 1990). To our knowledge, no direct investigation into the comparative response behavior of the cell line THP-1 has been carried out so far. In this study we hope to provide insights into the behavior of THP-1 if exposed to different human pathogens, and evaluate its ability to develop specific responses. To address how THP1 cells respond to the different stimuli, we investigated transcriptional profiles via RNA sequencing for the human THP1 cell line after exposure to the above mentioned pathogens and chemicals. To reduce analytical bias from the individual experiments, our analysis began with raw RNA sequencing data available at the NCBI Sequence Read Archive for the individual projects. A common, standardized pipeline was applied to carry out the data processing. We focused the subsequent analysis on the inter-project response for the individual pathogens. In order to overcome the very different experimental setups, we relied on more global approaches for tanscriptomic analysis. An important focus was the ability of the cancer derived THP-1 cell lines general response to distinguish the individual pathogens. Specific responses have been investigated in the individual experiments, yet such a variety of pathogens is expected to trigger substantially different overall response pathways. We investigated the impact of the individual pathogens via dimensionality reduction based clustering, and comparative GO term enrichment. Both on large scale to compare the overall behavior of the cells, and on the two available time course analyses to investigate the more minute temporal dynamic of transcription shifting. The available time course analyses comprise the bacterium and intracellular pathogen *Mycobacterium abscessus* and the yeast *Candida parapsilosis*. The *mycobacteria* species consist of a range of bacteria best studied for causing tuberculosis. They are documented to be fast growing and potentially multidrug resistant. The *M. abscessus* complex is also resistant to disinfectants and, therefore, can cause postsurgical and postprocedural infections (Lee et al., 2015). Candida species cause common nosochomial infections (Casadevall and Pirofski, 1999). As yeast, their mechanisms of pathogenicity differs significantly from that of bacteria. The yeast potentially inducing a much weaker response, testing the limits of THP-1 transcriptome adaptation.

## 2 Material and Methods

### 2.1 RNA sequencing data processing

RNA sequencing data was obtained in its raw read format from the Sequence read archive (Leinonen, Sugawara, and Shumway, 2011), with the exception of the data for *C. parapsilosis*, which was provided by the authors of Toth et al. (Tóth et al., 2017), the full list of sequence runs can be found in Table 1. A shell script to initiate the full download can be found at the github repository https://github.com/Gabaldonlab/THP1data. After extraction using the sratoolkit. Trimming, for quality pre-processing, of the reads was performed via Trimmomatic v0.32 (Bolger, Lohse, and Usadel, 2014). We mapped the reads using the STAR (Dobin et al., 2013) mapper against the hg38 human reference genome. Human genomic data hg38 v 81., both the reference and annotation files were obtained from UCSC (Ucsc and Browser, 2003). Reads were counted using the htseq package (Anders, Pyl, and Huber, 2015). For the analysis only annotated exons were considered. An overview of samples considered is presented in Table 1.

**Table 1:** Table of RNA seq runs processed in the analysis. The columns indicate, in this order: number of samples, pass of the mentioned quality check and accession number.

| Pathogen | Sample count | Time course | Quality control | Archive |
| --- | --- | --- | --- | --- |
| *Mycobacterium bovis* | 14 | No | failed | ERR5604 |
| *Mycobacterium abscessus* | 19 | Yes | pass | SRR23160 |
| Zika | 2 | No | pass | SRR51901 |
| Ebola | 2 | No | failed | SRR16602 |
| Marburgvirus | 2 | No | failed | SRR16367 |
| *Leishmania mexicana* | 4 | No | pass | SRR1562 |
| *Coxiella burnetii* | 4 | No | pass | SRR1562 |
| *Candida parapsilosis* | 16 | Yes | pass | from author |
| *Staphylococcus aureus* | 9 | No | failed | ERR50285 |
| Non Pathogen | 5 | No | pass | SRR16365 |

### 2.2 Data processing

Read count normalization was performed via transcript per million TPM normalization. R libraries were used to investigate the Principal Components underlying the data variability. The R built in prcomp module and the library FactoMineR (Le, Josse, and Husson, 2008) were used to compute the PCA and cluster estimation respectively. Tree based hierarchical clustering was carried out using the python scipy library. Gene enrichment was analyzed via python scripts available on the projects github https://github.com/Gabaldonlab/THP1data, generating a background model of variance. The enrichment compared to the full human background was carried out using the GOrilla tool (Eden et al., 2009)

### 2.3 Visualization

Visualization was performed via the R module ggbiplot, based on the ggplot2 library, as well as the FactoMineR and superheat plotting function for the respective R scripts. Visualization in python was produced via matplotlib.

### 2.4 Enrichment analysis

Expression enrichment for unregulated genes was computed for each gene on the variance over the non pathogen derived conditions, uninfected cells and separately against the chemicals ethanol and calcitriol. Outliers were tested against a normal distribution using student t-test. The method described by Benjamini & Hochberg (Benjamini and Hochberg, 1995) was used to correct for multiple testing and evaluate false discovery rate (FDR), in order to correct the resulting p-values. Adjusted p-values of < 0.05 were considered significant and analyzed by GOrilla against a total background.

## 3 Results and Discussion

As shown in Figure 1 the Principal Component Analysis showed a clear and distinct response to the individual conditions.

This can be considered an important sign that the THP1 cell line has retained its potential for detection of the individual pathogens in initiating individual responses. In a PCA, the abstract underlying effects are quantified and shown on components or axes, with relative strength per axis denoted in percent of variance explained. Individual principal components can show multidimensional response factors. Dimensions are visualized in Figure 1 Due to the complexity of analysis, the first four dimensions were considered to explain sufficient variance, collectively accounting for 36.5% of the observed variance. Similar responses were observed between the intracellular bacterial pathogens *Coxiella burnetii* and *Mycobacterium abscessus*, derived from experiments performed by independent groups [Table 1], suggesting that the first components are not influenced by the sequencing but directly by the THP-1 response, the profile of the two strains diverges in the third component showing more nuanced differences in response of THP-1 between the two pathogens.

**Figure 1:**
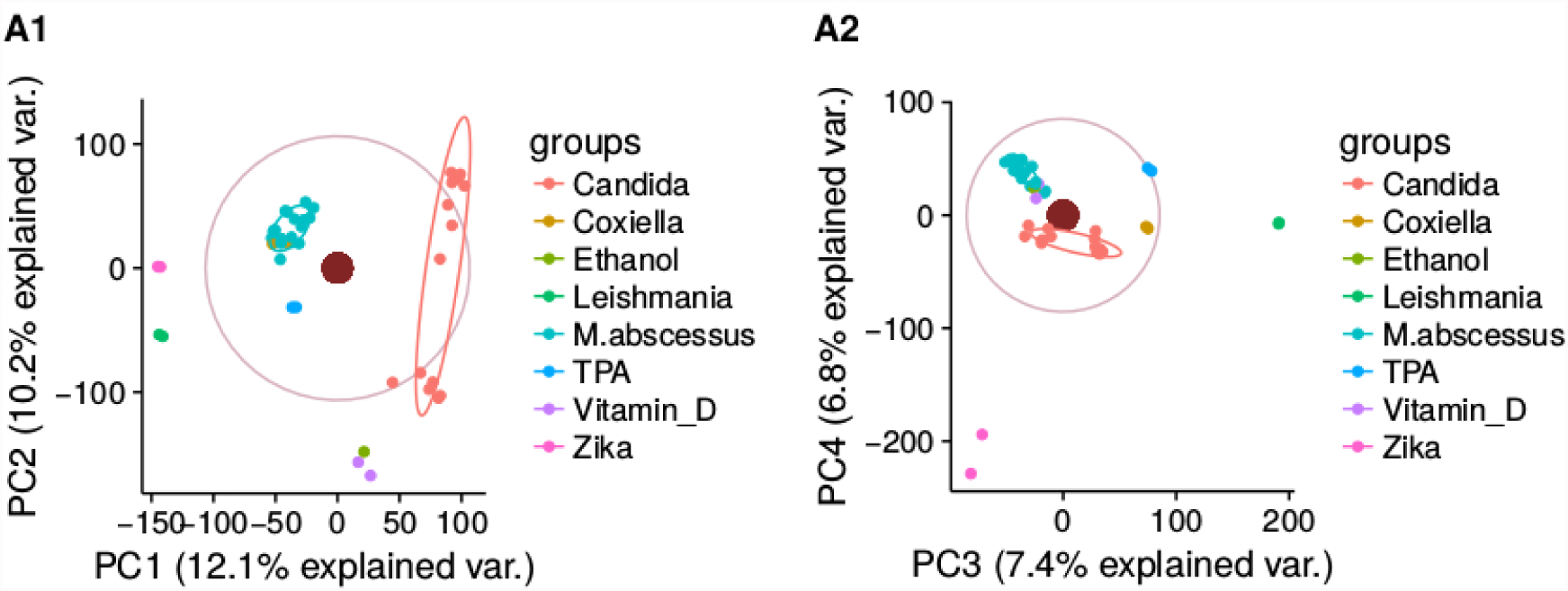
Principal component analysis for all genes, normalized to TPM. Due to the relative low variation per component the first four dimensions are displayed in two 2dimensional plots. Components are displayed in two plots PC1 and 2 in plot A1 and PC3 and 4 in A2. In A1, a cluster separation between the different pathogens and stressors can be observed on the first component. Two clusters comprised of the timeline experiments on *M. abscessus* and *C.parapsilosis* are visible along the first axis. A2 seperates viruses more clearly from the protist, as well as the intracellular bacteria *Coxiella bruneii* and *M. abscessus*.

Overall four distinct response clusters can be observed. With separation of clusters occurring for virus to yeast in the first component, and a distinction of bacteria over the second and third. The intracellular mycobacterium *M. abscessus* and the yeast *C. parapsilosis* are the most robust groups due to a larger sample size of 19 and 16 runs respectively, and replicates over a time course of infection assay. Factorial analysis to investigate time point responses was therefore limited to those two species. To gain a more detailed view on the minute behavior we investigated the response to the two larger time course analysis. Data was produced for rough and smooth morphologies during exposure times of 1, 4 and 24 hours. Although the analysis shows a clear separation (see Figure 2), the primary component derives from the effect between the replicates. The response to *C. parapsilosis* was less homogeneous, most likely due to the lack of replicates and the overall lower pathogenicity of *Candida* as compared to *Mycobacteria*.

**Figure 2:**
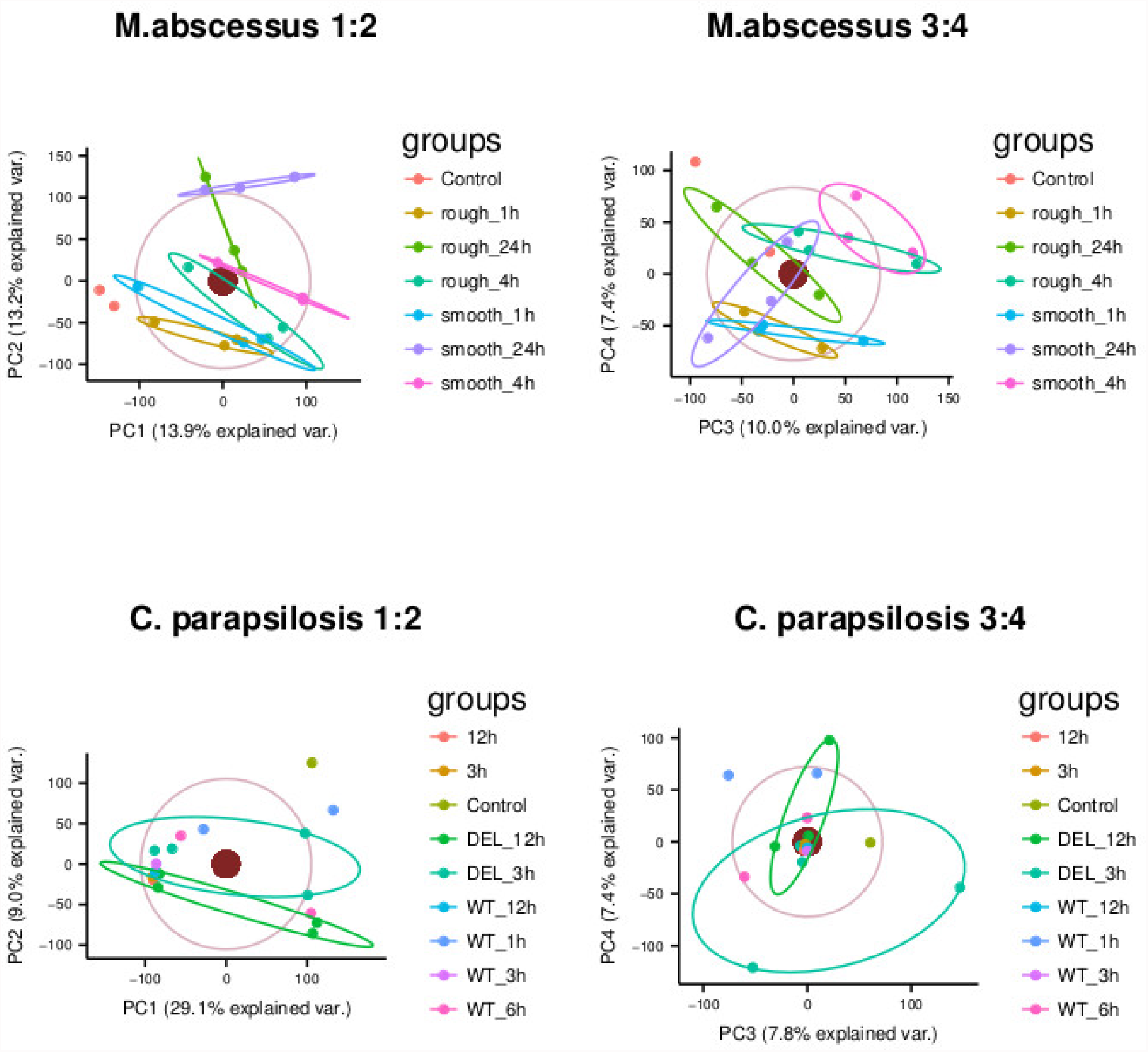
PCA of the time course analysis for the infividual sets of *M. abscessus* (above) and *C. parapsilosis* (below). Both studies show variance between technical replicates to be responsible for their first component, suggesting a lack of regulation by the THP-1 cells, or rather a lack of designated response. While *M. abscessus* shows a distinguishable variability over the time course, clustering on *C. parapsilosis* does not separate the individual conditions.

In the next step, we quantified overall transcriptional responses against background noise models visualized in Figure 3. Two Noise models were designed. In the first, (background) we used the average counts per gene in uninfected samples, to evaluate the expression compared to an uninfected baseline. For the second model (stressors) we evaluated the average gene count for samples treated with TPA, ethanol and the Vitamin D metabolite calcitriol to evaluate basic stress responses. Individual genes were tested for upregulation only against the model genes, returning a value of significance per gene and pathogen. This methods ignores experimental design in order to generalize the various experiments. Figure 3 shows the overlap between the background and stressor comparison. In total, 9302 genes were activated in any pathogen response compared to both backgrounds, with 1318 genes overlapping between the noise models. 6780 and 1204 genes were unique to the background and stressors model, respectively. This indicates an active response by THP-1, and its ability to distinguish uninfected surroundings to chemical stimuli.The cells show a clear distinction between the responses to the individual pathogens. Yet, the THP-1 response to *C. parapsilosis* shows no significant difference to the background model, but does present a distinction to the stressors model. *C. parapsilosis*, as a pathogenic yeast, is often commensal e.g (Gabaldon, Naranjo-Ortiz, and Marcet-Houben, 2016), and seemingly does not provoke a general strong transcriptomic response in the host. Interestingly, compared to the stressor background, only the *M. abscessus* cells show a classified defense response. According to GO terms, the response is antiviral [Table 2].

**Figure 3:**
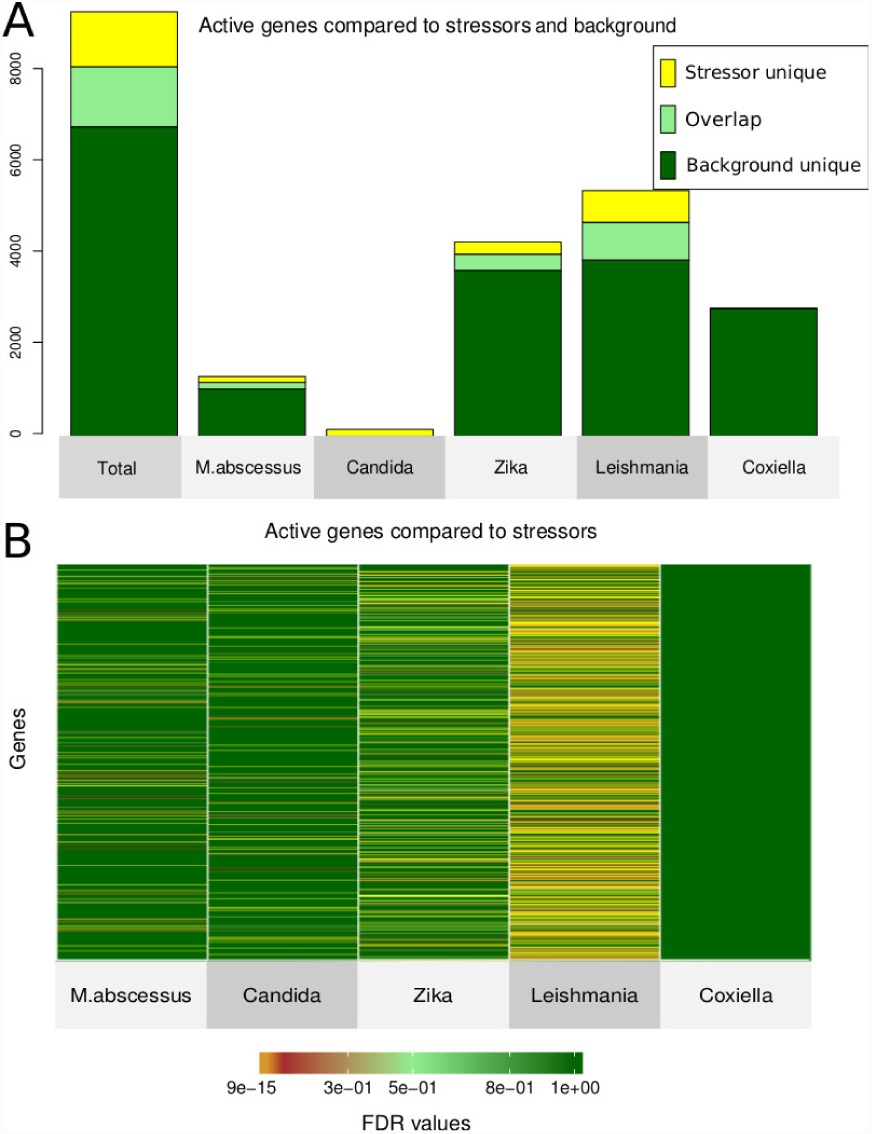
(A) Barplot visualizing the quantification of response against the two background models. Unique responses in yellow and dark green are only observed against the specific background. Overlap in light green shows responses similar between stress and uninfected cells. (B) shows a heatmap of individual genes actively regulated in the 5 species against the stress response. Notably, *Coxiella burnetii* shows no significantly enriched genes against the stressor subset.

**Table 2:** GO enrichment of pathogen responses against a stressor background

| *Mycobacterium abscessus* |  |  |
| --- | --- | --- |
| RNA biosynthetic/metabolic process | defense response to virus | type I interferon signaling pathway |
| *Zika virus* |  |  |
| forebrain development | triglyceride catabolic process | neutral lipid catabolic process |
| *Leishmania mexicana* |  |  |
| transmembrane transport | ion transport | organic acid transmembr. transport |
| *Coxiella brunetti* |  |  |
|  | None |  |
| *Candida parapsilosis* |  |  |
| mitotic cell cycle process | leukocyte activation | neutrophil activation |

An antiviral response to *M. abscessus* is partially expected, due to the related *M. tuberculosis* ability to trigger Interferon responses (Prabhakar et al., 2003), and Interferon production in general T-cell reponses (Belardelli and Gresser, 2010). *L. mexicana* shows the strongest overall response. With 3861 more genes activated than in the background, and a response of 694 unique genes to the *L. mexicana* stress compared to the stressor background. GO enrichment using GOrilla visualized in Table 2 shows unique responses for the pathogens. Most notably, the most enriched GO term for Zika is GO:0030900, pertaining to anatomical structure and forebrain development. This is in line with Zikas clinical symptoms, such as microcephalus as described in this review (Paixo et al., 2016). Enrichment for *L. mexicana* showed active genes involved in iron transport, an observation in line with Huynh et aliis (Huynh, Sacks, and Andrews, 2006) discovery of iron transporters being essential for parasitic reproduction. Active Iron transportation could therefore be an expected host response.

## 4 Conclusion

Our analysis suggests that the THP-1 cell line is capable of distinguishing various cellular stresses, and provide individual responses to various chemicals and pathogens. It accurately portraits gene enrichment for e.g Zika clinical symptoms. The large scale transcriptomic response is uniquely different for each analyzed experiment, and the cells show similar responses to the intracellular pathogens e.g *Coxiella burnetii* and *Mycobacterium abscessus*. Yet, the experimental resolution is less pronounced in more detailed experiments. Time course analysis using THP-1 showed a stronger variation between technical replicates than the actual experimental course. A recent study by Schurch et al (Schurch et al., 2016) estimates the number of true positives in RNA sequencing with 3 replicates to be between 20 and 40%. Using the first components as indicators of variance, we estimate that at least 12% of the total variance are attributed to technical variation. By using general probabiliy, we can estimate the true positives for triplicates to be between 13.6 and 27.26% using THP-1 cells ( according to (1 − (1 − *α*)*^n^*)). Therefore, especially for the investigation of pathogens with an expected mild response, additional replicates are strongly recommended.

